# Mitonuclear interactions produce extreme differences in response to redox stress

**DOI:** 10.1101/2022.02.10.479862

**Authors:** M. Florencia Camus, Vassilios Kotiadis, Hugh Carter, Enrique Rodriguez, Nick Lane

## Abstract

Incompatibilities between mitochondrial and nuclear genes can perturb respiration, biosynthesis, signalling and gene expression. Here we investigate whether mild mitonuclear incompatibilities alter response to redox stress induced by N-acetyl cysteine (NAC). We studied three *Drosophila melanogaster* lines with mitochondrial genomes either coevolved (WT) or mildly mismatched (BAR, COX) to an isogenic nuclear background. Responses to NAC varied substantially with mitonuclear genotype, sex, tissue and dose. NAC caused infertility and high mortality in some groups but not others. Using tissue-specific fluorespirometry, we show that NAC did not alter H_2_O_2_ flux but suppressed complex I-linked respiration in female flies, maintaining a reduced glutathione pool. High mortality in BAR females was associated with severe suppression of complex I-linked respiration, rising H_2_O_2_ flux in ovaries and oxidation of the glutathione pool. Our results suggest that redox stress is attenuated by suppression of complex-I linked respiration, to the point of death in some mitonuclear lines.

## Introduction

Mitochondria are the hub of both energy transduction and intermediary metabolism. Perturbations in cell respiration can affect ATP synthesis, redox balance and Krebs cycle flux, with downstream implications for biosynthesis ^1^, repair ^2^, signalling ^3^, gene expression and phenotype ^4–6^. Despite this central position in metabolism, mitochondria are uniquely vulnerable to perturbation, as the respiratory complexes are mosaic assemblies encoded by two genomes, the mitochondrial and nuclear, which have very different rates and modes of evolution ^7,8^. As a result, sexual reproduction generates new mitonuclear combinations in every individual, with the potential to affect all aspects of growth and development. Serious incompatibilities between mitochondrial and nuclear genes have been found in natural populations of several species, generated through introgression between divergent populations ^9–13^. These studies highlight the potential severity of mitonuclear incompatibilities between polymorphisms within normal populations, as opposed to pathological mutations known to cause mitochondrial diseases ^14,15^.

More subtle mitonuclear mismatches, stemming from single nucleotide polymorphisms (SNPs) can also profoundly affect gene expression and fitness ^16–18^, yet their metabolic basis remains to be elucidated. Crucially, the consequences of such subtle mitonuclear interactions might be amplified or attenuated by stress. To gain an insight into this question, we analysed the effect of mitonuclear interactions on redox stress in *Drosophila melanogaster*, a model that enables tight control over mitonuclear parameters. *D. melanogaster* has a mitonuclear system that is closely analogous to that of humans: mtDNA encodes the same core subunits of respiratory complexes, while the global divergence between populations, in terms of mitochondrial SNPs, is also similar ^19^. In this study, we therefore compared three different fly lines with subtle differences in the mitochondrial genome, all set against an isogenic nuclear background: WT (*w*^*1118*^); COX, whose mtDNA only differs from WT by one SNP in the gene for COXII (complex IV); and BAR, which has nine SNPs difference in protein-coding mtDNA, mostly in complex I and complex IV (**Fig. 1**).

**Fig. 1.**
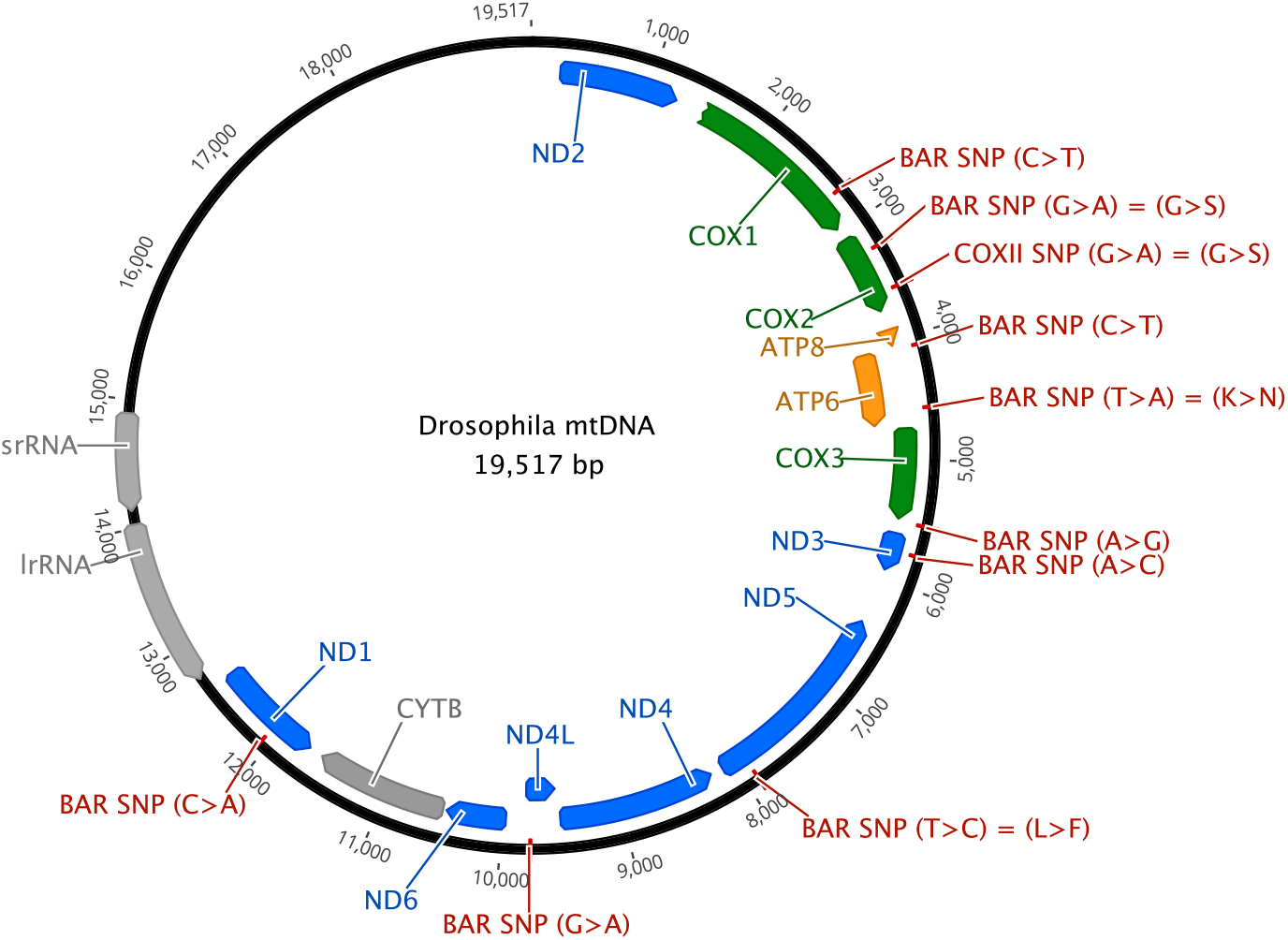
mtDNA genome showing single nucleotide polymorphism (SNP) differences between mitochondrial genotypes.

Because the three fly lines differ only in their mtDNA, the primary deficit must be in respiratory function or reactive oxygen species (ROS) flux ^20,21^. We therefore considered how the three mitonuclear genotypes altered metabolic and phenotypic responses to the glutathione agonist N-acetyl cysteine (NAC), which could potentially interfere with ROS flux, signalling and redox balance. Insofar as mitochondrial ROS signalling modulates mitochondrial biogenesis ^22^, and thus respiratory function, we anticipated that NAC might also modulate the respiratory rate. We considered low-dose (1 mg/mL) and high-dose (10 mg/mL) NAC, in line with earlier work in flies, which showed an increase in lifespan in males on 10 mg/mL NAC, but some toxicity at higher doses ^23^. More recent work showed toxicity with 10 mg/ml NAC ^24^, which we corroborate.

Here, we show that there can be large differences in fertility and survival between mitonuclear genotypes in males and females depending on NAC dose. We used tissue-specific high-resolution fluorespirometry to measure multiple respiratory parameters linked in real-time with H_2_O_2_ flux as an indication of overall ROS flux. Glutathione redox state generally remained stable except in the BAR females, which had high mortality associated with severe suppression of complex I-linked respiration, rising ROS flux in ovaries, and oxidation of the glutathione pool, suggesting terminal loss of redox control. Our study demonstrates that mitonuclear interactions can indeed determine individual response to redox stress, with very different outcomes in male versus female flies.

## Results

### Mitonuclear genotype and sex can have major fitness effects in response to NAC

Both BAR and COX flies have been reported to exhibit mild male subfertility at 25 °C, which in the case of COX is exacerbated by higher temperatures (29 °C) ^25,26^. We confirmed this effect here: both male genotypes showed a slight decrease in the fertility of young adults (day 12) at 25 °C (**Fig. 2A**). We found a significant interaction between mitochondrial genotype and NAC treatment for each sex (female: F = 31.205, p = < 0.01; male: F = 18.193, p < 0.001). Low-dose NAC (1 mg/mL: NAC 1) had little or no effect on either male or female fertility, but high-dose NAC (10 mg/mL: NAC 10) had a severe effect on the fertility of females in all three fly lines, lowering the number of offspring by 60 to 90 %. In contrast, while NAC 10 lowered male fertility in WT and COX flies, it had no overall effect on the fertility of BAR males.

**Fig. 2.**
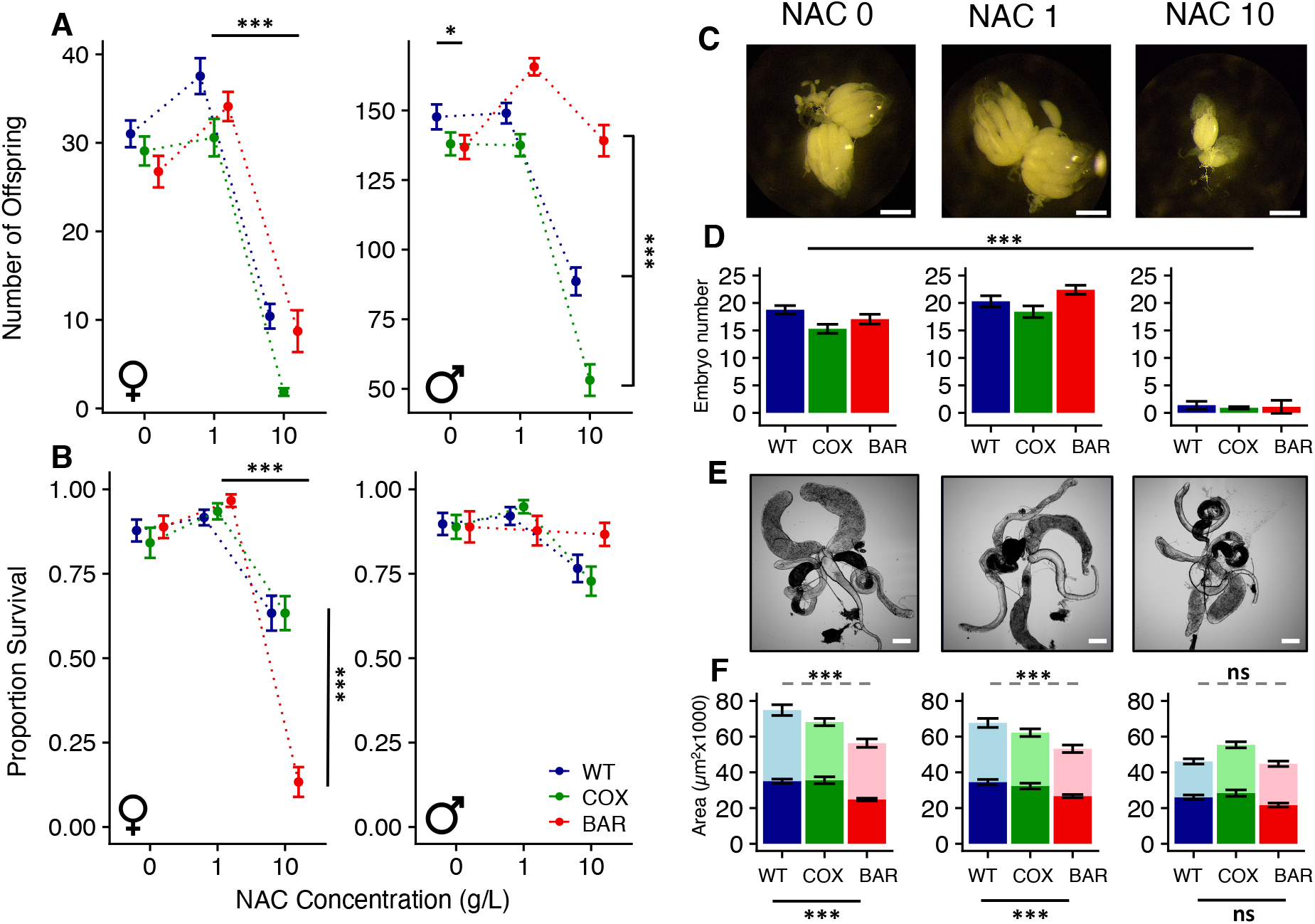
Components of fitness for all mitochondrial genotypes and NAC treatments. (**A**) Total number of offspring produced (mean ± SE) for males and females after a single mating event across all mitochondrial genotypes and NAC treatments. (**B**) Proportion of adult flies exposed to the three NAC treatments that survived to the fitness assay (day 12). (**C**) Images of WT ovaries after 12 days of being exposed to the three NAC treatments (representative of all 3 genotypes). (**D**) Quantification (mean ± SE) of ovariole number across all mitochondrial genotypes and NAC treatments. (**E**) Images of WT male reproductive tissue (testes, accessory glands, seminal vesicle) after 12 days exposure to the three NAC treatments. (**F**) Male reproductive tissue surface area (μm^2^ x 1000) for all mitochondrial genotypes across all NAC treatments (mean ± SE). Dark bars are testes surface area, whereas light bars are accessory gland measurements.

During the setup of the fitness experiment, we recorded how many flies survived NAC exposure, that is, how many flies died before measuring their fitness (**Fig. 2B**). Whereas NAC 1 had little effect on the survival of either male or female flies in any of the mitonuclear lines, NAC 10 clearly affected the survival of female flies (NAC: χ^2^ = 118.862, p < 0.001). In particular, BAR females were particularly affected by NAC 10, with barely 1 in 10 surviving to day 12 (female - mito × NAC: χ^2^ = 13.442, p = 0.009, **Fig. 2B**). In contrast, while NAC 10 did depress overall male survival (NAC: χ^2^ = 20.7268, p < 0.001), BAR males were unaffected. Thus, the effects of NAC 10 on survival at day 12 were sexually antagonistic in BAR flies: females mostly died, while males were barely affected by NAC 10. This pattern of sexual antagonism was not apparent in either WT or COX flies, which have fewer SNPs difference in mtDNA.

The differential fitness effects of high-dose NAC were also clear at the level of sexual organs. NAC 10 shrank the ovaries of all three fly lines (representative examples in **Fig. 2C**), matched by a collapse in embryo number (NAC: F = 215.8642, p < 0.001, **Fig. 2D**). Males showed a more nuanced pattern (**Fig. 2E and F**). Untreated BAR males had smaller testes (WT_0_-BAR_0_: p < 0.001, COX_0_-BAR_0_: p < 0.001) and accessory glands (WT_0_-BAR_0_: p = 0.0298) than WT at day 12 (**Fig. 2F**), which corresponded to their subfertility (**Fig. 2B**), indicating a specific mitonuclear incompatibility in the testes. Treatment with NAC 10 shrank the testes and accessory glands of both WT and COX flies but had little further effect on BAR males (Testes-NAC: F = 17.9905, p < 0.001; Acc-NAC: F = 17.6898, p < 0.001, **Fig. 2F**), consistent with their uncompromised fertility at day 12 on NAC 10 (**Fig. 2B**).

### Differences in longevity can be sexually antagonistic

Our analysis of these components of fitness indicates that fly lines with isogenic nuclear background but distinct mtDNA are affected differently by NAC at day 12. We asked whether these differences in fitness persisted across the life course. Without treatment, BAR females lived significantly longer than either WT or COX females (WT-BAR: χ^2^ = 89.491, p <0.001; COX-BAR: χ^2^ = 89.912, p <0.001, **Fig. 3**). NAC 1 had small effects on longevity, depending on the mitochondrial genotype. The lifespan of WT females was unchanged on NAC 1, while COX flies had a shorter lifespan (COX_0_-COX_1_: χ^2^ = 42.144, p < 0.001) and BAR females had a slight increase (BAR_0_-BAR_1_: χ^2^ = 6.076, p = 0.009). NAC 10 was again severely toxic to BAR females (BAR_0_-BAR_10_: χ^2^ = 203.099, p < 0.001), which were nearly all dead before day 20. However, WT and COX females were also seriously affected just a few days later (WT_0_-WT_10_: χ^2^ = 213.729, p < 0.001; COX_0_-COX_10_: χ^2^ = 167.755, p < 0.001, **Fig. 3**). Thus, while BAR flies were significantly worse hit by NAC, the severe toxicity of NAC 10 was not specific to BAR females (mito × NAC: χ^2^ = 172.9350, p < 0.001).

**Fig. 3.**
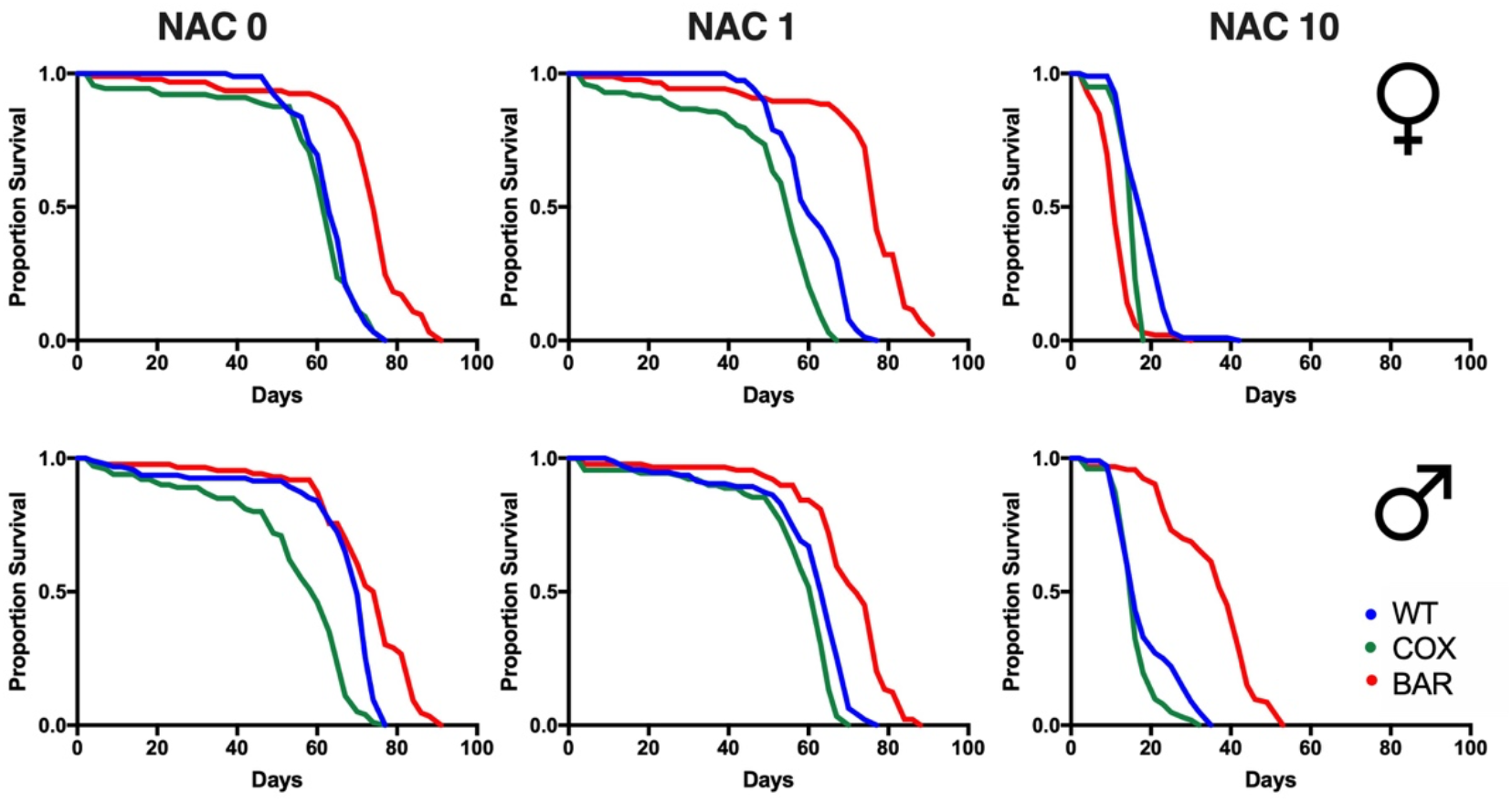
Survival plots for all mitochondrial genotypes and NAC treatments.

Males showed significant effects of both mitochondrial genotype and NAC on longevity (mito × NAC: χ^2^ = 172.9350, p < 0.001, **Fig. 3**). Without treatment, COX flies were significantly shorter lived than either WT or BAR males (p < 0.05). NAC 1 abolished the difference between WT and COX flies, with both groups now displaying similar longevity (**Fig. 3**). In contrast, BAR males had significantly longer lifespan than either of those groups (*p* < 0.05). As in females, NAC 10 again had severe toxicity in all groups, albeit less devastating for males (**Fig. 3**). In this case, far from being the worst hit, BAR males had relatively long lifespan (WT_10_-BAR_10_: χ^2^ = 67.398, p < 0.001; COX_10_-BAR_10_: χ^2^ = 159.288, p < 0.001), again exhibiting signatures of sexual antagonism.

### Metabolic profile and effect of NAC vary by tissue

We employed tissue-specific high-resolution fluorespirometry to simultaneously measure respiratory rates and H_2_O_2_ flux for two tissues (thorax and reproductive) in each sex. We chose to first inspect for broad patterns using Principal Components Analysis (PCA), with results showing distinct metabolic profiles for different tissue (**Fig. 4;** for loadings see **Fig. S1 and Table S1**). Male and female thoraces exhibited overlapping metabolic profiles, whereas reproductive organs diverged from the thoraces. From the PCA loadings we know that much of the variance captured by PC1 is related to oxygen metabolism, whereas variance for PC2 is driven by H_2_O_2_ flux. Based on this, we can infer that ovaries had much lower metabolic rates and lower stress, while testes had similar metabolic profile to ovaries, but significantly more stress. While it is possible that we over-estimated metabolic stress by homogenizing the testes and accessory organs compared with permeabilizing muscle fibres, we homogenized ovaries using the same methodology, yet H_2_O_2_ flux was > 10-fold greater relative to oxygen consumption in the testes compared with the ovaries (i.e. ROS flux ratio, **Fig. S2**). Clearly NAC 10 significantly affected the ovaries, showing a major increase in stress, possibly indicating a pro-oxidant effect through reductive stress. Other shifts with NAC did not achieve statistical significance in the pooled data, but oxygen metabolism was suppressed in female versus male thoraces; in fact, the PCA plots masked specific differences in oxygen metabolism between mitonuclear lines.

**Fig. 4.**
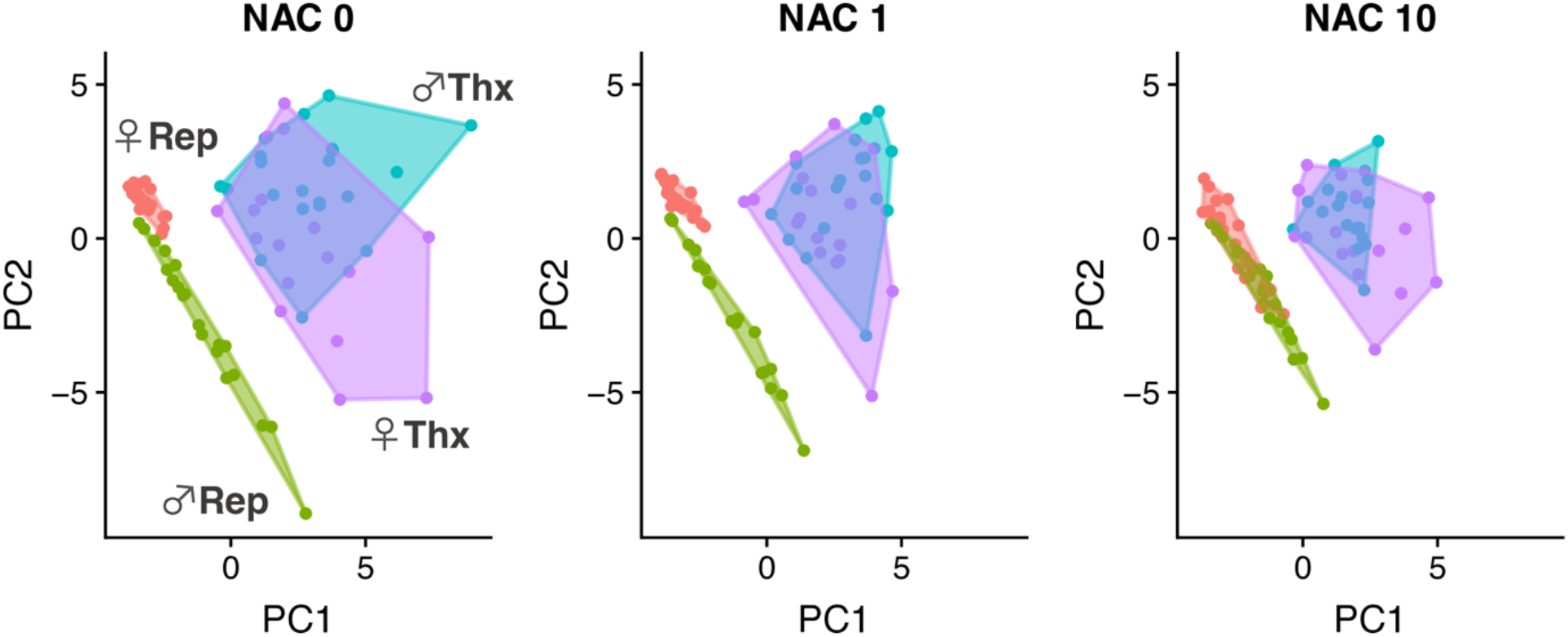
Principal component analysis by tissue and NAC treatment. Tissue-specific respirometry results for all mitochondrial genotypes combined. Respirometry data was partitioned using principal components analysis across all NAC treatments. The first two principal components account for most of the variance (PC1 = 54.03% and PC2 = 36.52%). For loadings see Fig. S1.

### NAC suppresses complex I-linked respiration in vulnerable female lines

We next examined specific metabolic states across all samples (sex/nac/genotype combinations). State 3 respiration (**Fig. 5**) is initiated when ADP is added to pyruvate, malate and proline, and so gives an indication of respiration linked with substrates that feed electrons into the respiratory chain mainly through complex I. Because complex I-linked respiration maximises proton pumping and so ATP yield, complex I-linked substrates tend to be utilized most when aerobic demands are high (as in the thorax here). Full traces for all respiratory states are shown in **Fig. S3**. We found that state 3 respiration is around one third higher in females than in males (F = 26.6778, p < 0.001, **Fig. 5A and B**), corroborating earlier work ^27,28^. While NAC 1 had little effect on respiratory rates, NAC 10 decreased state 3 respiratory flux in pooled female lines (F = 4.7432, p = 0.011; **Fig. S4**). The suppression of state 3 flux varied significantly with mitonuclear genotype (F = 3.2287, p = 0.04423). Untreated COX and BAR females had slightly lower state 3 respiration than WT, albeit still substantially higher than any of the male fly lines. While respiration was lowered across all female thoraces with NAC 10, it was severely depressed in BAR females (WT-BAR: p = 0.0402, COX-BAR: p = 0.0046, **Fig. 5A and B**) at day 12 – the same group whose survival was cut by NAC 10 treatment.

**Fig. 5.**
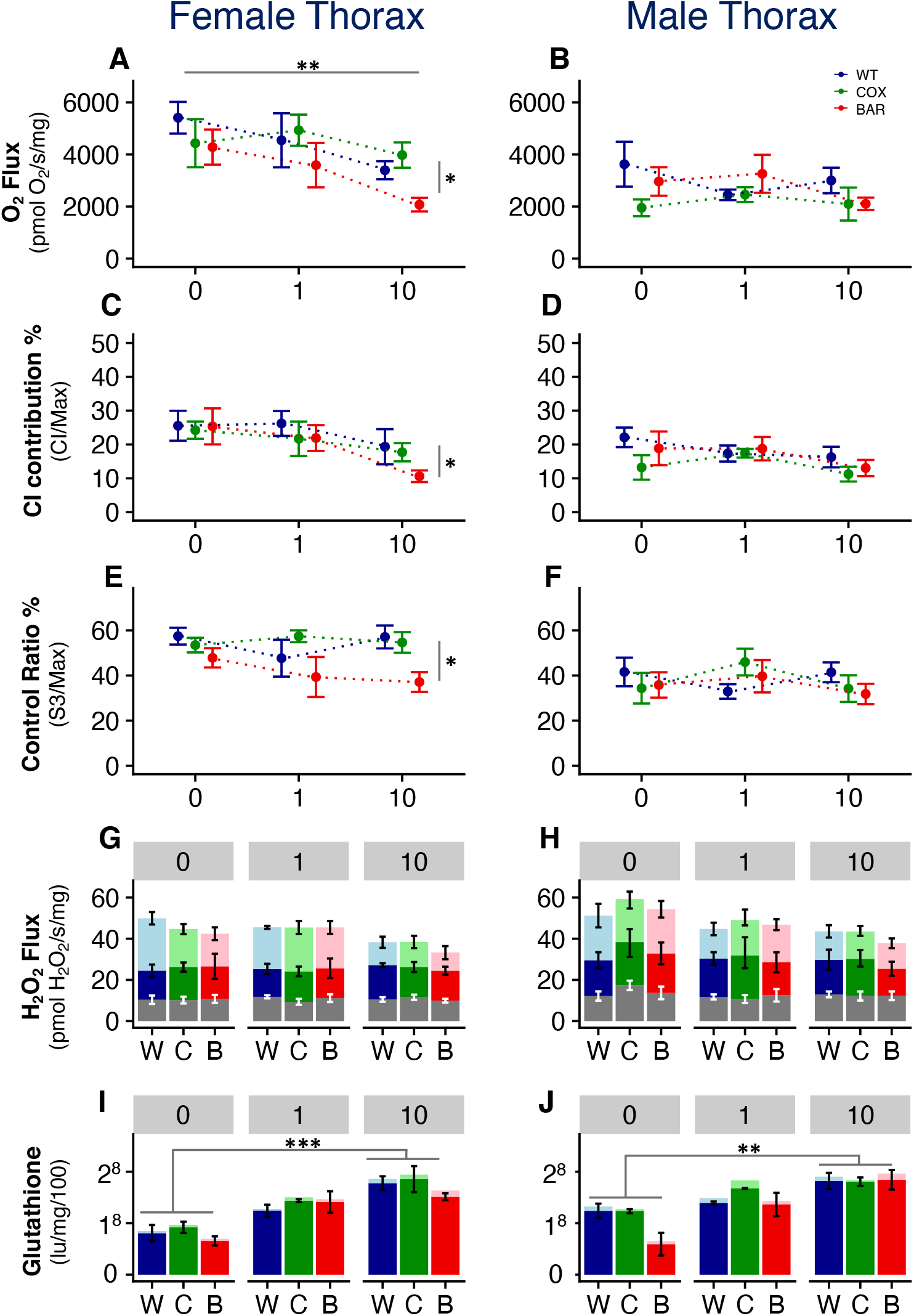
Oxygen flux, H_2_O_2_ flux and redox balance in thoraces. Fluorespirometry and glutathione results (mean ± SE) for female (left) and male (right) thoraces for all mitochondrial genotypes and NAC treatments. (**A and B**) State 3 O_2_ flux; (**C and D**) complex I contribution; (**E and F**) control ratio; (**G and H**) H_2_O_2_ flux; (**I and J**) glutathione concentration. For H_2_O_2_ flux, grey bars denote state 3 flux; dark-coloured bars denote flux after addition of rotenone; and light-coloured bars show flux after addition of antimycin A. For glutathione samples, dark-coloured bars show reduced glutathione and light-coloured bars oxidised glutathione, with the combined coloured area being total GSH.

Maximum coupled respiration was also significantly depressed by NAC 10 across all female genotypes in the thorax (NAC × sex: F = 3.3013 p = 0.04134, **Fig. S5**), but in this case to similar absolute values across all genotypes. Because survival at day 12 was ~15 % for BAR females compared with ~60% for WT (**Fig. 2**), it is unlikely that the suppression of maximum respiration was responsible for the death of BAR females. This inference is reinforced by two analyses of relative respiratory rates, both of which indicate specific suppression of complex I (**Fig. 5C and D**). First, complex I contribution to maximum respiration (based on rotenone inhibition) roughly halved from ~25 % for WT and BAR females when untreated (NAC 0) to ~12 % of maximal respiration for BAR females on NAC 10, compared with ~20 % for WT and COX females (**Fig. 5C and D**). Second, the control ratio (state 3 respiration / maximum coupled respiration) fell from around 50–60 % in all untreated female lines, to less than 40 % in BAR females on NAC 10, while remaining above ~55 % in WT and COX females on NAC 10 (**Fig. 5E and F**). These findings are consistent with the hypothesis that NAC 10 suppressed state 3 respiration, and specifically complex I activity, in the thoraces of BAR females at day 12, while sparing WT and COX flies at this same time point. The differences between complex I activity and control ratio are most likely explained by the use of proline as one of the three substrates for state 3 respiration, as this can also feed electrons directly into complex III via proline dehydrogenase ^29^.

While the thoracic state 3 respiration in males was little more than half that of males, NAC 10 suppressed pooled state 3 respiration to a much lesser extent (NAC: F = 4.7432, p = 0.011, **Fig. 5 and Fig. S4**). In fact, on a per mg protein basis, the absolute rate of state 3 respiration was similar in BAR males and females on NAC 10; the difference lay in the degree of suppression relative to untreated controls (~20 % for males versus ~50 % for females). Despite their lower state 3 respiration, male mortality at day 12 was limited in comparison with females. In contrast, the difference in maximum respiration was less pronounced, at around 15 %, suggesting that male flies are better able to survive on complex III-linked substrates such as glycerol phosphate (**Fig. S5**). Furthermore, none of the male groups showed significant depression in maximum coupled respiration on NAC 10 (**Fig. S5**). This tendency for male flies to depend less on complex I-linked substrates ^27^ was confirmed by complex I activity and control ratios (**Fig. 5D and F**). CI activity was ~20 % lower, and control ratio ~30 % lower than females, but these values were not significantly depressed by NAC 10 in any fly line. Taken together, these results suggest that male flies have less need for complex I-linked respiration than females, and this lower dependence may partially protect males against NAC toxicity.

### Respiratory suppression at complex I maintains redox balance in the thorax

Given the major differences in respiratory control between males and females, and between different mitonuclear lines, H_2_O_2_ flux remained remarkably stable in the thoraces of both male and female flies (**Fig. 5G and H, and Fig. S6)**. Total glutathione levels increased significantly with higher doses of NAC across all tissues (NAC_0_-NAC_1_: p < 0.001, NAC_0_-NAC_10_: p < 0.001). This confirms that the flies did indeed ingest NAC with their food, and that it had the anticipated effect of raising tissue glutathione levels, with NAC 10 raising thoracic glutathione levels more than NAC 1 (NAC_1_-NAC_10_, p < 0.001, **Fig. 5 I and J**). Despite this sustained increase in glutathione levels in both male and female thoraces, state 3 H_2_O_2_ flux generally remained stable, while as noted previously, respiration was suppressed, mainly at complex I, and especially in BAR females.

Despite the substantial rise in thoracic glutathione levels with NAC, the ratio of oxidised to reduced glutathione barely changed with increasing NAC dose, suggesting that modulation of ROS flux prevented the oxidation of the glutathione pool, maintaining redox homeostasis in the thorax (**Fig. S7**). The effect of respiratory inhibitors suggests that ROS flux was indeed modulated by suppression of respiration. The stacked bars of **Fig. 5G and H** show H_2_O_2_ flux after inhibition of maximum respiration at complex I, using rotenone, and at complex III, using antimycin A. The stacked bars reflect cumulative H_2_O_2_ flux, i.e. ROS flux increased on addition of rotenone and again on addition of antimycin A. These measurements demonstrate that the H_2_O_2_ flux derives from mitochondrial complexes I and III rather than from elsewhere in the cell. Most importantly, H_2_O_2_ flux changed very little in response to rotenone in the thoraces of both male and female flies of any mitonuclear line, and at any dose of NAC. This implies that the maximum reactivity of complex I to oxygen remained stable across all these states. That this low reactivity was achieved by limiting the number of reactive FeS centres in complex I was shown by a corresponding suppression of complex I activity and state 3 respiration on NAC 10. Most ROS are thought to derive from complex I under physiological conditions ^30–32^, hence state 3 respiration is preferentially suppressed. This interpretation is borne out by ROS flux ratios (**Fig. S2**), which show little to no change in maximal H_2_O_2_ flux per unit oxygen consumption in the maximal coupled respiratory state (max H_2_O_2_ / max coupled respiration), reflecting the most physiological input of electrons from multiple substrates simultaneously.

### NAC disrupts redox balance in sex organs

The tight control of redox balance in the thoraces of flies was lost in the sexual organs with NAC 10. In terms of overall dynamics, respiratory control in the ovaries reflected that in the thoraces, albeit at ~10-fold lower flux levels (**Fig. 6 and Fig. S4**). This difference may have partly reflected tissue damage from homogenization. While the different preparations meant that direct comparisons should not be made between flux rates in the thoraces and sex organs, the patterns of respiratory control were reliable, and could be compared between ovaries and testes prepared using the same methods. These again showed sexually antagonistic responses in the reproductive tissues (NAC × sex: F = 11.7907, p < 0.001, **Fig. S5**). In the ovaries, NAC treatment suppressed both state 3 and maximum respiration (**Fig. 6; Fig. S4 and S5**). Unlike the thorax, however, maximum respiration was more strongly reduced than state 3 respiration. In contrast, in male reproductive tissue, NAC 10 tended to increase both state 3 and maximum respiration (**Fig. 6; Fig. S4 and S5**). H_2_O_2_ flux was less tightly constrained in the reproductive tissue, increasing with NAC in the ovaries, and significantly so in BAR flies on NAC 10, while falling slightly in testes and accessory glands on NAC 10 (NAC × sex: F =6.2105, p = 0.002, **Fig. 6**). Tissue-specific glutathione levels rose on NAC 10 in all cases, and dramatically so in BAR females, where glutathione levels rose around 8-fold compared to the other genotypes (**Fig. 6**). Despite these shifts, the ratio of oxidised to reduced glutathione remained stable in the ovaries, and tended to become more oxidised in the testes and accessory glands (**Fig. S7**). However, the suggestion that glutathione pool became more oxidized in BAR females in both thoraces and ovaries was borne out by pooling data from both organs (**Fig. S8**). In this case, the glutathione pool was significantly more oxidized in BAR females, indicating redox stress in this group alone – the group dying at day 12.

**Fig. 6.**
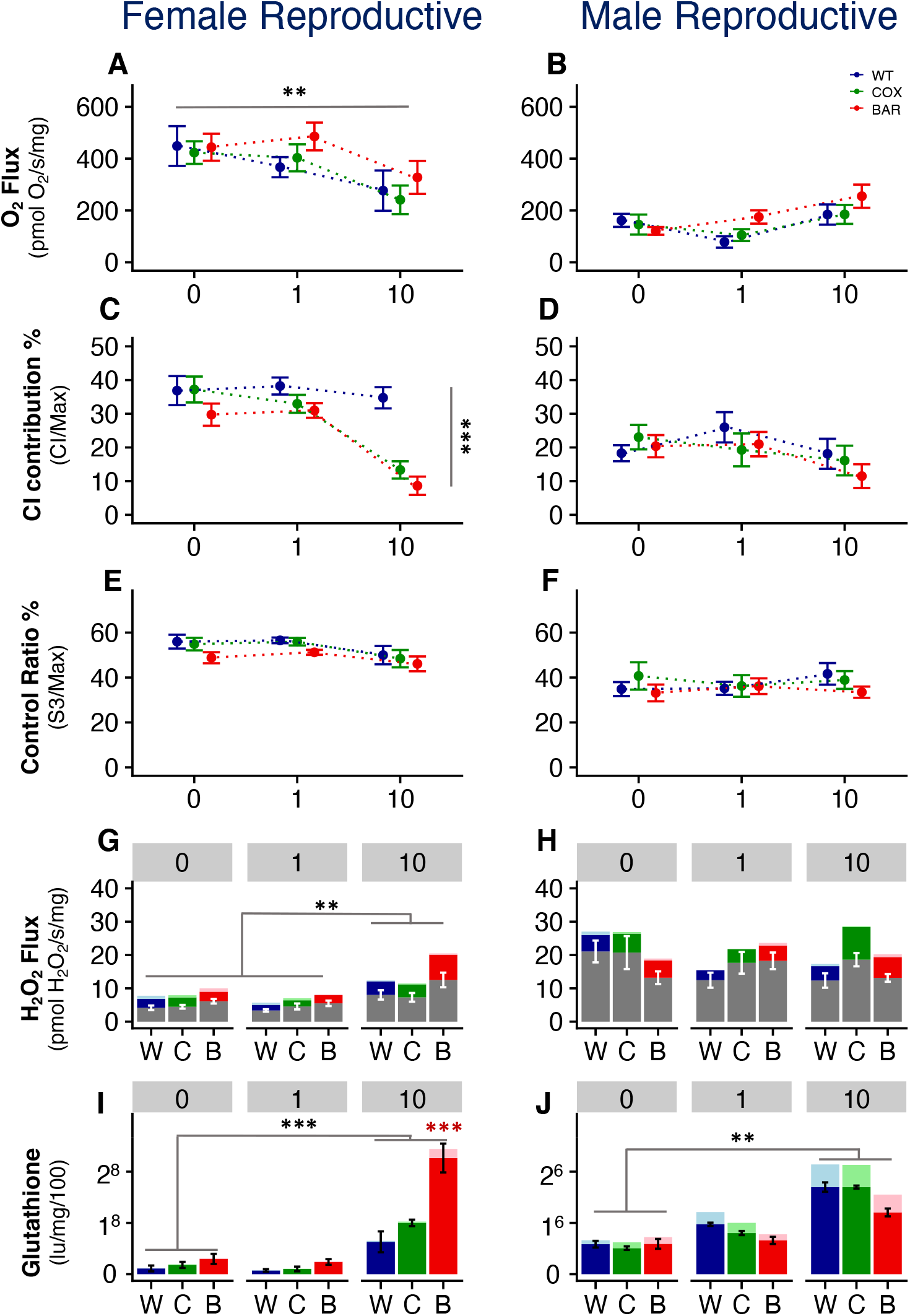
Oxygen flux, H_2_O_2_ flux and redox balance in reproductive tissues. Fluorespirometry and glutathione results (mean ± SE) for female (left) and male (right) reproductive tissues, for all mitochondrial genotypes and NAC treatments. (**A and B**) State 3 O2 flux; (**C and D**) complex I contribution; (**E and F**) control ratio; (**G and H**) H_2_O_2_ flux; (**I and J**) glutathione concentration. For H_2_O_2_ flux, grey bars denote state 3 flux; dark-coloured bars denote flux after addition of rotenone; and light-coloured bars show flux after addition of antimycin A. For glutathione samples, dark-coloured bars show reduced glutathione and light-coloured bars oxidised glutathione, with the combined coloured area being total GSH.

The main reason why ovaries tended to lose redox balance again appears to be a dependence on complex I-linked respiration. Complex I contribution in the ovaries was a quarter higher than the thoraces, at around 40 % in all three mitonuclear lines (**Fig. 6C**). Likewise, the control ratio was 50–60 % in the ovaries, as high as thoraces (and around a third higher than testes), again implying a high demand for ATP synthesis through complex I-linked respiration, most probably reflecting the high metabolic costs of egg production ^33,34^ (**Fig. 6E**). NAC 10 strongly suppressed complex I contribution in both COX and BAR flies, to around 25 % of maximum respiration, with the control ratio falling to 40–50 % (**Fig. 6C**). Notably, BAR females on NAC 10 had significantly elevated H_2_O_2_ flux in normal state 3 respiration (p = 0.0472, **Fig. 6G**) and after rotenone inhibition (p = 0.0326, **Fig. 6G**). Presumably, suppression of complex I activity was not sufficient restrict ROS flux. This failure to retain redox balance through respiratory modulation was also indicated by a sharp rise in the ROS flux ratio (max H_2_O_2_ / max O2) in the ovaries in all three mitonuclear lines on NAC 10 (NAC × sex: F = 12.3134, p < 0.001, **Fig. S2**).

H_2_O_2_ flux in the male reproductive tissues is difficult to decipher, being high in almost all respiratory states, especially given the very low rates of respiration in the male sex organs – around 20-fold lower than the male thoraces. While these values might in part reflect tissue damage during homogenization, they also probably reflect specialization of mitochondria to processes other than ATP synthesis, which is known to contribute relatively little in sperm ^35^ and stem cells ^36^. For example, ROS may drive cell growth and proliferation through transcription factors such as HIF_1α_ ^37^. This interpretation is supported by the relatively low H_2_O_2_ flux in the reproductive tissues of untreated BAR males (NAC 0; **Fig. 6H**), which corresponded to significant male subfertility (**Fig. 2A**). NAC 1 slightly increased H_2_O_2_ flux in BAR males (**Fig. 6H**), as well as state 3 (**Fig. 6B**) and maximal coupled (**Fig. S5**) respiration, all of which were linked with significantly improved fertility (**Fig. 2A**). While NAC 10 reversed many of these metabolic effects in BAR males (**Fig. 6 and Fig. S5**) the total glutathione pool was restricted in size more successfully than in other fly lines (**Fig. 6**), corresponding to virtually unimpaired fertility in young adults (**Fig. 2B**). There were no differences in the oxidation state of the glutathione pool in either the testes (**Fig. S7**) or whole BAR flies (**Fig. S8**).

## Discussion

Subtle differences in mitonuclear interactions are generated between individuals by sexual reproduction. Despite their potential to perturb energy metabolism, biosynthesis, signalling and gene expression, mitonuclear interactions have been neglected in genome-wide association studies of disease risk or drug responses ^38^. Here, we have used a *Drosophila* model to probe the effect of subtle mitonuclear interactions on responses to N-acetyl cysteine (NAC), which is usually considered to be an antioxidant that promotes glutathione synthesis ^39^. In untreated flies, mitonuclear interactions had relatively subtle phenotypic effects, notably mild male subfertility (**Fig. 2**) and modest differences in lifespan (**Fig. 3**). These subtle mitonuclear interactions underpinned major differences in drug responses to high-dose NAC (10 mg/mL). In general, NAC 10 did not alter H_2_O_2_ flux (**Fig. 5**) but suppressed complex I-linked respiration in female flies more than male flies (**Fig. S4**), while in most cases maintaining a reduced glutathione pool (**Fig. S8**). However, BAR flies, which differ from wild-type in just 9 SNPs in protein-coding mtDNA (**Fig. 1**) showed dramatic sexually antagonistic responses to NAC. Male flies were not affected by NAC in either survival or fertility at day 12 (**Fig. 2**) but most females had died by then, with the few survivors also showing severely compromised fertility (**Fig. 2**). The high mortality rate in BAR females was associated with severe suppression of complex I-linked respiration (> 50%) in both thoraces (**Fig. 5**) and ovaries (**Fig. 6**), rising H_2_O_2_ flux in ovaries (**Fig. 6**) and oxidation of the glutathione pool in pooled tissues (**Fig. S8**), indicating loss of redox control.

Our results suggest that suppression of respiration at complex I normally acts as a flux capacitor to maintain redox balance. Unlike immediate regulation of ROS flux (which we do not address here) long-term redox balance is governed by S-glutathionylation of proteins, which is known to suppress complex I-linked respiration in response to oxidation of the glutathione pool ^40^. The suppression of respiration at complex I lowers ROS flux from reactive FeS clusters, maintaining redox balance in the glutathione pool, but at the cost of aerobic capacity, as shown in **Figs. 5** and **6**. We will address the mechanisms by which complex I activity is modulated in future work. The hypothesis that suppression of complex I maintains normal redox balance is supported here by the observations that (i) H_2_O_2_ flux was similar in state 3 respiration and after rotenone inhibition, which reflects the architectural capacity of complex I to generate ROS (**Figs. 5** and **6**); (ii) the ROS-flux ratio (H_2_O_2_ flux relative to maximal coupled respiration; **Fig. S2**) remained stable, which reflects most closely physiologically balanced respiration on multiple substrates, including those known to increase ROS flux, such as succinate and glycerol phosphate; and (iii) the redox state of the glutathione pool barely changed, except in the ovaries of BAR females on NAC 10 (**Fig S7** and **S8**), where the maximal suppression of complex I-linked respiration failed to mitigate redox stress (**Fig. 6**).

The suppression of complex I is both more pronounced and more costly in female flies, where the high metabolic costs of egg production ^33,34^ likely demands greater dependence on complex I. In general, female flies have slightly greater maximal respiratory rates (**Fig. S4**) but nearly double the rate of state 3 respiration in both thoraces and ovaries, implying greater complex I-dependence than males; conversely, males have greater rates of complex III-linked respiration, linked with the substrate glycerol phosphate. Complex I-linked respiration maximises oxidation of mitochondrial NADH, promoting both ATP synthesis and Krebs-cycle flux, which provides many precursors for biosynthesis needed for egg production. If female fitness requires greater complex I activity, this presumably places physiological limitations on how much complex I can be suppressed in female flies, which in turn limits how effectively ROS flux can be suppressed. Maximal suppression of complex I in BAR females on NAC 10 can therefore explain the rise in H_2_O_2_ flux in the ovaries. The fact that NAC did not act as an antioxidant but instead induced reductive stress is supported by the clear oxidation of the glutathione pool in NAC females alone (**Fig. S8**). We infer from this that increased ROS flux from heavily reduced respiratory complexes was compensated by down-regulation of complex I, suppressing state 3 respiration to the limits where redox control was lost, which likely caused the early death of female BAR flies (**Fig. 2**).

While our interpretation of respiratory suppression and redox balance has not been proved here (ongoing transcriptomic and metabolomic work will address these questions) there is no doubt that subtle differences in mitonuclear genotype can induce extreme differences in responses to redox stress through NAC supplementation. This raises two important questions. First, to what extent can our findings be generalised to other drug responses? And second, how relevant are inbred mitonuclear fly lines more generally, and in particular to human populations? In terms of generalisability, our results show that compensation for stressed mitonuclear incompatibilities primarily involves complex I. This complex is increasingly recognized as a critical hub in most age-related diseases, from Alzheimer’s disease to cancer ^41^, and yet the contribution of any mitonuclear interactions to disease risk has barely been considered ^42–46^. Given that many interventions also target complex I, even if downstream, we anticipate that other drug responses will be attenuated or amplified by mitonuclear interactions. Indeed, we and others have already shown substantial differences between mitonuclear genotypes in response to high-protein diets, so a systematic consideration of mitonuclear interactions in drug responses is overdue ^47–51^.

In terms of extension to human populations, the fly model utilized here enables tight control over mitonuclear variables, which allows us to demonstrate clear effects against an isogenic nuclear background. Most people differ from each other in at least as many SNPs in mtDNA as the fly lines reported here, but in addition of course vary in nuclear alleles, which would be expected to modulate risk. We know from mitochondrial diseases that nuclear variations can indeed suppress or unmask severe symptoms ^52^, and we have found similar effects in a large panel of fly lines in which 9 different mtDNAs were set against 9 different nuclear backgrounds (81 mitonuclear genotypes) ^53^. But the extent to which ordinary SNPs in both mitochondrial and nuclear genes influence life-long health, risk of disease and drug responses is simply unknown. The iceberg of missing hereditability has long confounded understanding of genetic risk factors from genome-wide association studies ^54^, yet the relative risk associated with specific nuclear SNPs should vary in unpredictable ways depending on their interactions with mitochondrial SNPs, whether direct or indirect. Mitochondrial SNPs are often neglected on the grounds that there are few of them compared with nuclear SNPs, but their importance to ATP synthesis, redox balance, biosynthesis, signalling and gene expression mean that they have a greater potential to amplify risk ^38^. Our results here demonstrate that subtle mitonuclear interactions not normally associated with overt disease can cause unexpected differences in physiological response to redox stress. Such effects are likely to be pervasive, and we urge that mitonuclear interactions are considered as the basis of a fresh approach to personalised medicine, which might be termed pharmacomitogenomics.

## Materials and Methods

### *D. melanogaster* maintenance and strains

All *Drosophila melanogaster* stock strains were maintained on a standard mix of molasses/cornmeal medium at a constant 25°C on a 12/12 light dark cycle. Three different strains of *D. melanogaster* were used in this experiment, differing only in their mitochondrial genomes. The “WT” strain is the coevolved strain, with the w^1118–5095^ nuclear genome naturally coevolved with the mitochondrial genome. The second strain has the same isogenic w^1118–5095^ nuclear background but has a mitochondrial haplotype termed “COXII”. The COXII haplotype is derived from w^1118–5095^ flies in which the mitochondrial mutation COII^G177S^ has become fixed ^25^. COII^G177S^ is a single non-synonymous change to subunit II of cytochrome C oxidase, and this SNP is the only difference between COXII and WT mtDNA (Figure 1). The third strain has a mitochondrial haplotype designated BAR, and is derived from a wild population from Barcelona, Spain. For this strain, the original chromosomes have been replaced by those of the w^1118–5095^ nuclear genome through the use of a balancer chromosome crossing scheme ^55^. Mitochondrial DNA from BAR flies differs from WT mtDNA by 9 SNPs, mostly in protein-coding genes ^56^ (Figure 1). All fly strains are kept in a strict breeding regime, whereby female flies from all strains are consistently backcrossed to the isogenic w^1118–5095^ nuclear background, which itself is propagated via full-sub crosses. This regime ensures that all fly strains maintain the nuclear background as similar as possible.

For the purposes of the fitness experiment, competitor flies were used to mate with our focal (mitochondrial) flies. These competitor flies were from the outbred *D. melanogaster* genotype L_HM_ ^57^ and reared in the exact same conditions as experimental mitochondrial flies. Outbred flies were used to ensure that the focal flies were breeding with competitors of the highest possible fitness so all fertility responses were attributable to the focal flies alone.

### NAC media preparation

For conditions requiring the antioxidant N-Acetyl-L-cysteine (NAC - Sigma A7250), fly food media was prepared at concentrations of 1mg/ml and 10mg/ml. For this, the desired quantity of NAC was dissolved in 100ml of water and added to 900ml of liquid fly food media. Once the NAC solution and liquid media were thoroughly mixed, 4mL of NAC food were dispensed into individual fly vials. Powdered NAC and media stocks were stored at 4°C and warmed to room temperature before use.

### Fertility assay and survival to reproduction estimates

For these experiments males and females were assayed separately but following the same experimental rearing regime. Newly eclosed virgin *D. melanogaster* adults from the BAR, COXII or WT strains were immobilised through exposure to low concentrations of CO_2_ (5ppm), split by sex and transferred to vials containing treatment media at 0 (Control), 1 or 10 mg/ml NAC. Each vial contained three males or three females from one strain and each combination of media/strain had forty replicates per group (N= ~120 flies).

Flies were maintained on their respective treatment media for twelve days and transferred to new treatment media three times a week. After twelve days on treatment media, flies were transferred to vials containing standard control media and which had three competitor flies of the opposite sex to mate with. Flies were transferred to new media after two days and then adults discarded after a further two days. Eggs lain on media during each two-day period were allowed to develop at a constant 25°C for fourteen days after which fertility was determined by a count of all adult progeny present in each vial.

Survival was measured as the percentage of each *D. melanogaster* strain for each media/sex combination that had survived by the final (16^th^) day of the fertility assay. Fly survival was recorded every time flies were swapped onto new food during the experiment.

### *Drosophila* reproductive tissue size

For male reproductive tissue measurements, 12-day old males corresponding to each mtDNA genotype and NAC treatment were dissected (total of 40 males per experimental treatment). For each individual, reproductive organs were extracted and placed on a microscope slide containing one drop of room temperature PBS solution. A cover slip was placed on top of the sample and an image was acquired using a Leica DMLB microscope. Surface area of testes and accessory glands for each image was acquired using ImageJ software.

Female ovaries were extracted individual flies of each experimental treatment when flies were 12 days of age. Ovaries were placed on a microscope slide containing a drop of PBS, and number of ovarioles was counted under a light microscope.

### Longevity

Newly eclosed *D. melanogaster* adults from the BAR, COXII or WT strains were collected within a 24 our period from each density-controlled vial. Vials for each haplotype were combined in a small holding cage with a petri dish of molasses-cornmeal-yeast food and left for 48 hours to mate. Following this 48-hour period, flies were immobilised through exposure to low concentrations of CO2 (5ppm), split by sex and transferred to vials containing treatment media at 0 (Control), 1 or 10 mg/ml NAC. Each vial contained ten males or ten females from one strain and each combination of media/strain had ten replicates per group (N= ~100 flies). Flies were transferred to fresh vials of their corresponding NAC treatment, and their survival scored three times a week.

### High-resolution Respirometry

High-Resolution Respirometry with Fluorometry (HHR-Fluo) was conducted with an Oroboros Oxygraph-2K (O2K). Calibration for O2 sensors and for H_2_O_2_ detection via Amplex Ultra Red according to manufacturer’s protocol. Protocols can be found online at www.bioblast.at. Thoracic muscle was permeabilised in ice cold BIOPS (CaK_2_EGTA 2.77 mM, K_2_EGTA7.23 mM, MgCl_2_.6H_2_0 6.56 mM, Imidazole 20 mM, Taurine 20mM, Na_2_Phosphocreatine 15mM, Dithiothreitol 0.5 mM, K-MES 50 mM, Na_2_ATP 5.77mM) containing 81.25 ug/mL Saponin for 30 minutes, with a subsequent 10 minute wash in ice cold MiR05 buffer. Testes and ovaries were homogenised in *Drosophila* Ringer’s buffer. Oxygen flux recordings were conducted in MiR05 ((pH 7.1) EGTA 0.5 mM, MgCl_2_.6H_2_O 3 mM, Lactobionic Acid 60 mM, Taurine 20 mM, KH_2_PO_4_ 10 mM, HEPES 20 mM, D-Sucrose 110 mM, BSA (essentially fatty acid free) 1 g/L) with the additions of Pyruvate (10 mM), Malate (10 mM), and L-Proline (10 mM), as well as HRP (2 units) and Amplex Ultra Red (10 μM) prior to insertion of sample. Oxygen Flux rates were measured before and after additions of ADP (5 mM) to observe State 3 respiration, Succinate (10 mM) to reach State 3 for CI+CII, snGlycerol-3-phosphate (10 mM) to achieve state 3 for CI+CII+G3PDH, and FCCP was titrated (0.25 μM increments) until the sample reached maximum uncoupled State 3 (State 3u). Once State 3u was reached the sequential addition of the inhibitors Rotenone (0.5uM) and Antimycin A (2.5 uM) allowed the estimation of CI and CII+ G3PDH relative contributions to overall flux. O_2_ and H_2_O_2_ flux rates were normalised to total protein (mg) in chamber by conducting Pierce™ BCA total protein quantification assay (Thermo Fisher Scientific) either directly from homogenate sample, or by homogenising thoraces retrieved from the O2K recording chambers. For each experimental unit (mito/tissue/treatment/sex combination) we collected between 6-8 replicates.

### Glutathione assay

Intracellular reduced and total glutathione levels were quantified using the bioluminescent Promega GSH-Glo™ glutathione assay (Promega), as recommended by the supplier. In brief, 5 thoraces, 40 ovaries and 60 testes were extracted from flies of each experimental treatment (in triplicate). Individual samples were first dissected and held in ice-cold Drosophila Ringers Buffer (1x) while dissections were underway. Samples were then spun down at 8000xG for 5 minutes at 4°C. Supernatant was removed and replaced with 100μl of DPBS+0.2mM of EDTA. Samples were then homogenised in this buffer, and later centrifuged at 17000xG for 10 minutes (4°C). Supernatant was transferred to a new microtube and stored at −80°C until time of assay.

For the assay, 15ul of each experimental sample were used to measure reduced glutathione and total glutathione separately on a 384-well plate (Greiner). For total glutathione measurements, 1mM of TCEP was added to each well and incubated at RT for 30 minutes. To all samples, 15ul of 2X GSH-GloTM Reagent was added and samples were incubated for 30 minutes at RT. Following this incubation time, 30ul of Luciferin detection reagent was added to each well, mixed well using a shaker and incubated for 15 minutes. Luminescence was then measured using a Pherastar FS. A standard was run alongside all samples which included 5, 2.5, 1.25, 0.625, 0.3125,0.15625,0.078125 and 0mg of glutathione. All samples were normalised to total protein concentration (mg) in the sample by using the Pierce™ BCA total protein quantification assay (Thermo Fisher Scientific).

### Statistical analyses

Male and female phenotypic traits were analysed separately, as experiments were performed separately. For female fitness, the overall total number of offspring was zero-inflated. We therefore analysed this dataset using a negative binomial distribution ^58^, in which the zero values are a blend of sampling and structural effects (negative binomial parameter; variance = ϕμ). These models were performed using the R (v. 3.0.2) package glmmADMB (http://glmmadmb.r-forge.r-project.org/glmmADMB.html). The response variable was total number of offspring produced, with mitochondrial haplotype and NAC treatment (plus their interaction) as fixed factors. Male offspring production was modelled by fitting a generalized linear mixed model, using a Gaussian distribution. For each analysis, mitochondrial strain and NAC treatment (plus their interaction) were modelled as fixed factors in the *lme4* package ^59^ in R ^60^.

Proportion survival to reproduction was modelled as a binomial vector, composed of total number of flies in the vial at the start of the experiment and number of flies that died, using a binomial distribution and logit link. Mitochondrial haplotype and NAC treatment (plus their interaction) were modelled as fixed factors. Analyses were performed using the *lme4* package ^59^ in R ^60^.

Reproductive morphological traits were analysed in two different ways. Male reproductive tissue surface area was modelled using a linear model, whereby area was a response variable, with mitochondrial genotype and NAC treatment (plus their interaction) as fixed effects. Female ovariole numbers were modelled using a general linear model with a quasipoisson distribution (to account for overdispersion of data). Ovariole number was the response variable, with mitochondrial genotype and NAC treatment (plus their interaction) as fixed effects. Models were implemented using the *lme4* package ^59^ in R ^60^.

Longevity analyses were performed using the *survival* package in R ^61,62^. We used Cox proportional hazard models, with survival as a response variable. Mitochondrial haplotype, NAC treatment and sex (plus all their interactions) were fixed effects in the model. We also investigated this dataset in a sex-specific manner. For this, we split the data into each sex respectively and ran models that included mtDNA and NAC treatment (plus their interactions) as fixed factors.

Metabolic continuous variables were centred and standardised prior analysis using the ‘prcomp’ function in R version 3.3.2 ^62^. Most of the respiratory parameters were highly correlated to each other, therefore we used principal component analysis to reduce these variables to two linearly uncorrelated components. The two main principal components accounted for 96.78% of the variance. Following the principal components analysis, we tested the significance of sex and tissue on PC1 and PC2. The main model had (residual) PC1 and PC2 as response variables, with tissue, sex and their interaction as fixed effects. We performed a Tukeys post-hoc test to determine which factors (sex and/or tissue) contributed to the overall multivariate effect. All analyses were performed using the *aov* and *TukeyHSD* functions in R version 3.3.2 ^62^ For subsequent metabolic analyses, we focused on key parameters that reflect mitochondrial respiration; State 3 oxygen, State 3 peroxide flux and glutathione. For these three traits we used linear models with either variable as response variables and sex, mitochondrial genotype and NAC as fixed factors (plus all their interactions).

Further analyses included using Tukeys post-hoc test to identify which factors were contributing to the overall multivariate effect. Analyses were performed using the *lmer* and *TukeyHSD* functions in R version 3.3.2 ^62^.

## Acknowledgments

**General**: We thank Rebecca Finlay for help with running experiments. **Funding:** This work was supported by funding from the Biotechnology and Biological Sciences Research Council (BB/S003681/1) and bgc3 to NL and Leverhulme Trust (RPG-2019-109) to MFC. **Author contributions:** All authors contributed to the conception and writing of the paper. MFC carried out the phenotypic work. VK, HC & ER carried out the fluorespirometry. All authors analyzed the results. MFC, ER and NL wrote the MS. **Competing interests:** NL is a member of the SAB of Metro International Biotech. **Data and materials availability:** All data is accessible on Dryad repository

