## Supplementary Data for "Mitonuclear interactions produce extreme differences in response to redox stress"

### Mitonuclear interactions cause extreme differences in drug responses

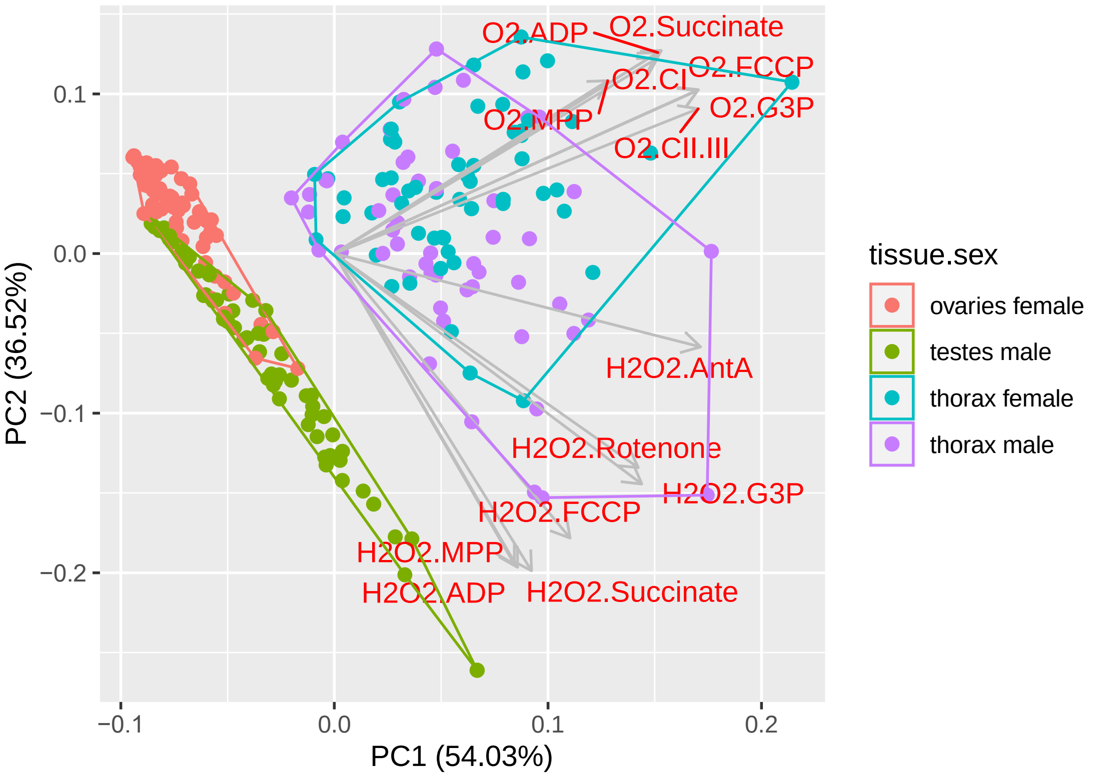


**Fig. S1 Principal component analysis for combined NAC treatments.**
Datapoints are coloured by tissue, with principal component loadings being displayed by the grey arrows. From the loadings, we conclude that PC1 is mainly driven by oxygen flux, whereas PC2 is driven by $H_2O_2$ flux.

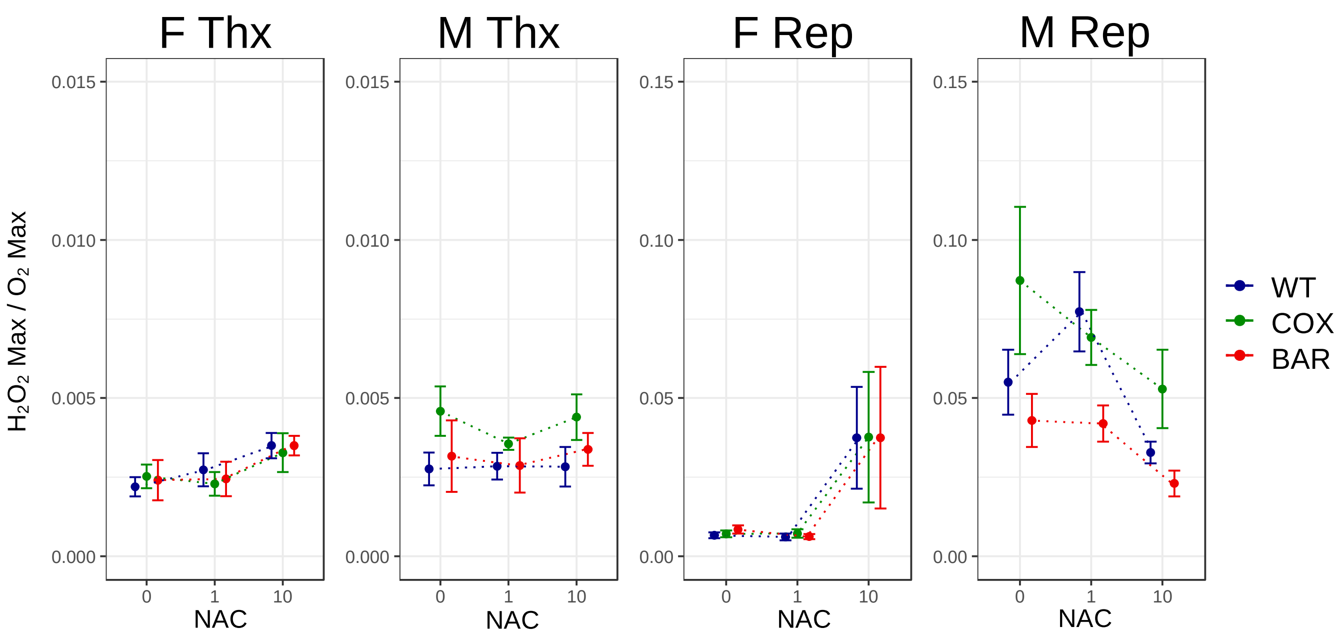


**Fig. S2 ROS flux ratios**

$H_2O_2$ flux during maximum coupled respiration/oxygen flux during maximum coupled respiration for all tissue/treatment combinations.

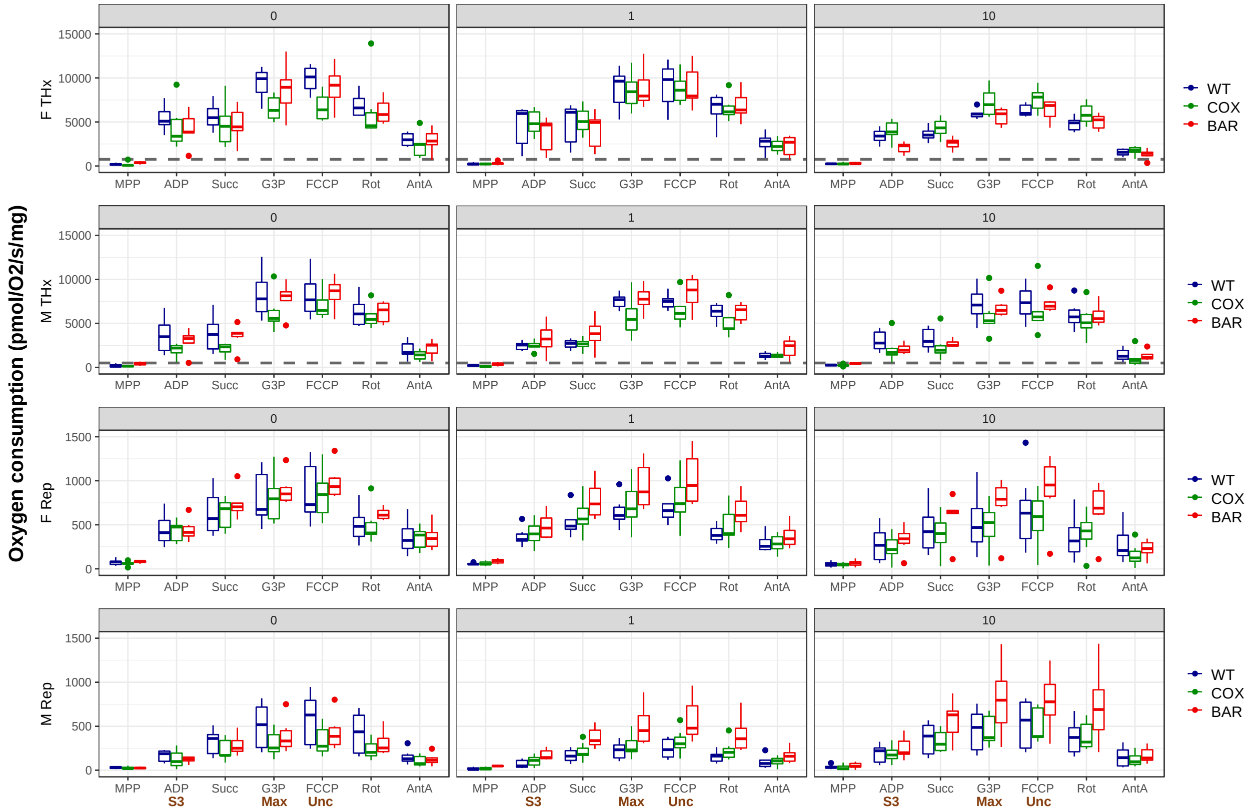


**Fig. S3 Oxygen flux as a function of the respiratory states for all tissues and NAC treatments.**

For the full description of respiratory states, see Methods section. State 3 (S3), maximum respiration (Max) and uncoupled respiration (Unc) are highlighted at the base of the plot for reference. Note the different scales for thorax and reproductive tissues; the dashed lines in the thorax plots indicate the average flux for the reproductive tissue of the same sex.

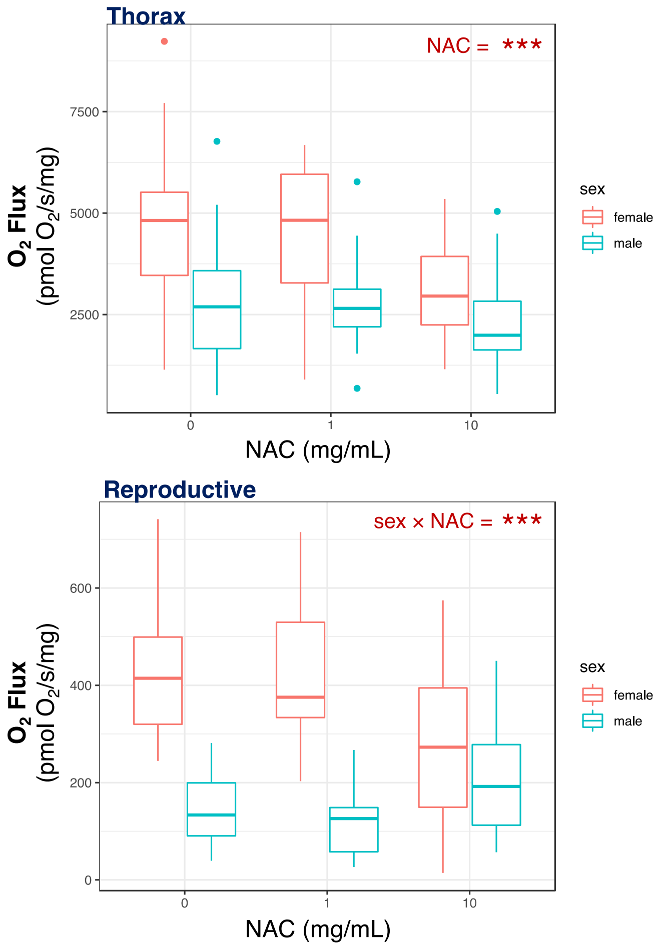


**Fig. S4 Changes in state 3 respiration with NAC for combined genotypes.** State 3 respiration was initiated with complex I substrates malate, pyruvate and proline. These plots combine all mtDNA genotypes to depict overall patterns for both sexes and tissues as a function of NAC concentration. See Table S2 for full ANOVA tables.

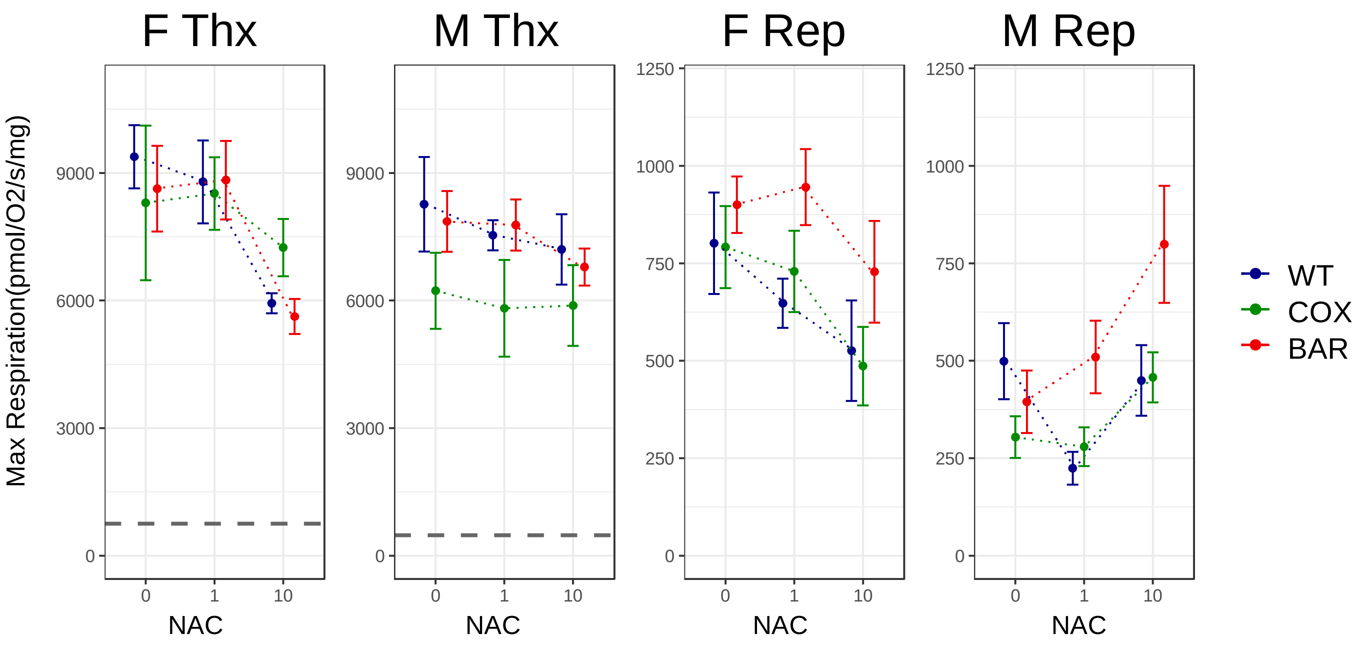


**Fig. S5 Maximum coupled respiration for all tissue/treatment combinations.** Maximum coupled respiration: ADP + complex I substrates = malate, pyruvate and proline; complex II substrate = succinate; and complex III substrate = glycerol-3-phosphate. Note the different scales for thorax and reproductive tissues; the dashed lines in the thorax plots indicate the average flux for the reproductive tissue of the same sex.

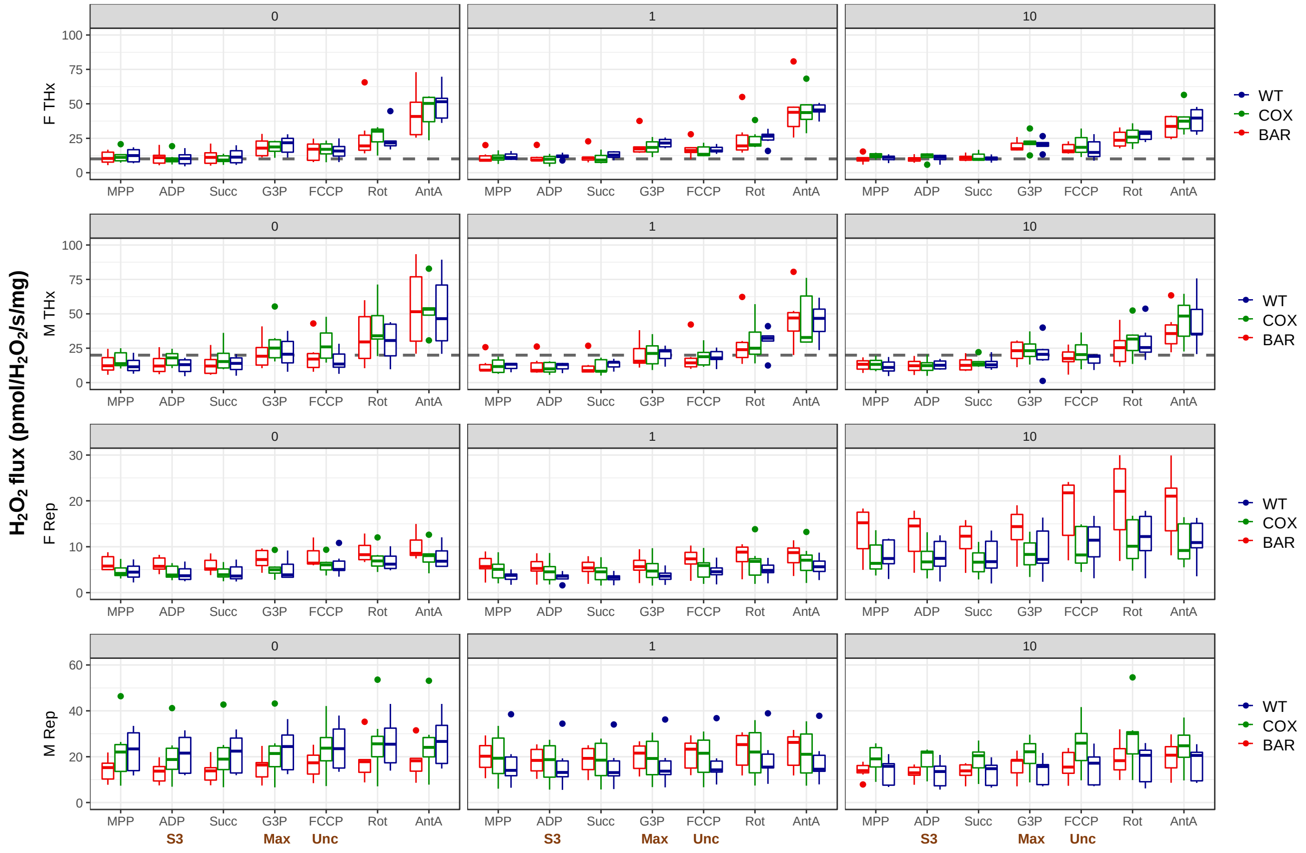


**Fig. S6 ROS flux as a function of the respiratory states for all tissues and NAC treatments.** Simultaneous real-time $H_2O_2$ detection via Amplex Ultra Red in all respiratory states. For the full description of respiratory states, see Methods section. State 3 (S3), maximum respiration (Max) and uncoupled respiration (Unc) are highlighted at the base of the plot for reference. Note the different scales for thorax and reproductive tissues; the dashed lines in the thorax plots indicate the average flux for the reproductive tissue of the same sex.

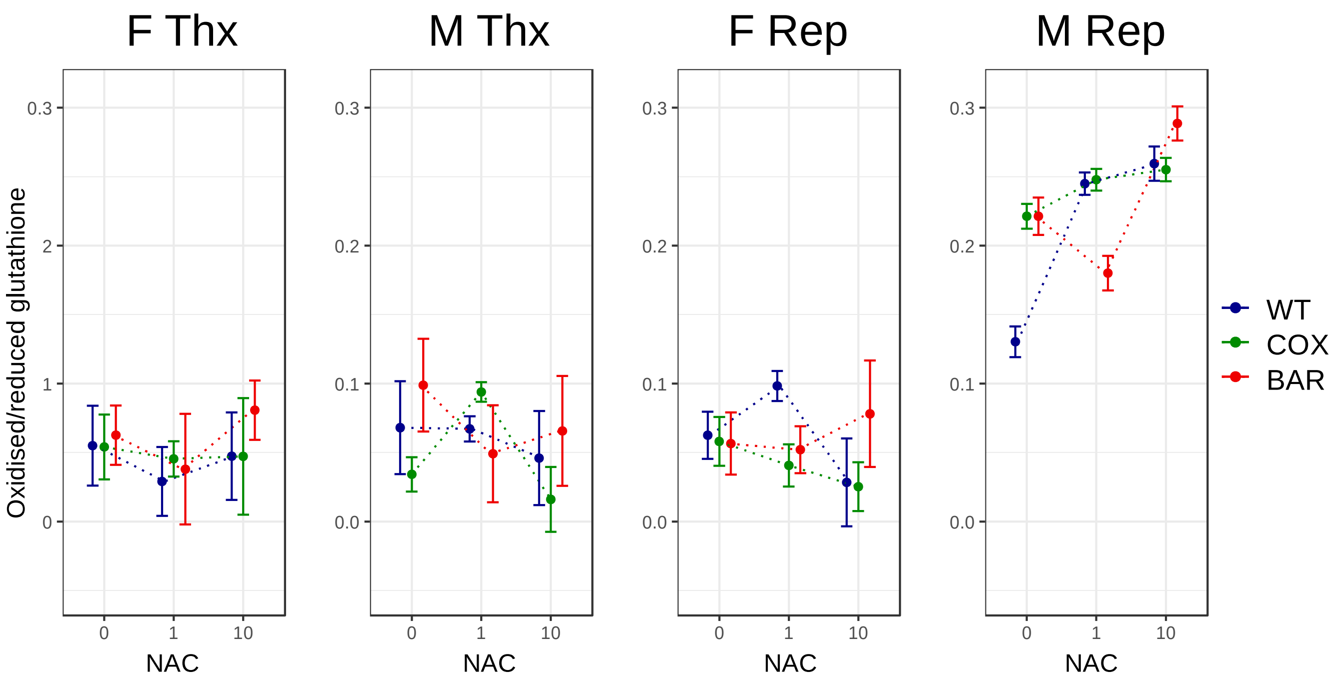


**Fig. S7  Glutathione redox state**
Ratio of oxidised to reduced glutathione across all tissue and mtDNA combinations.

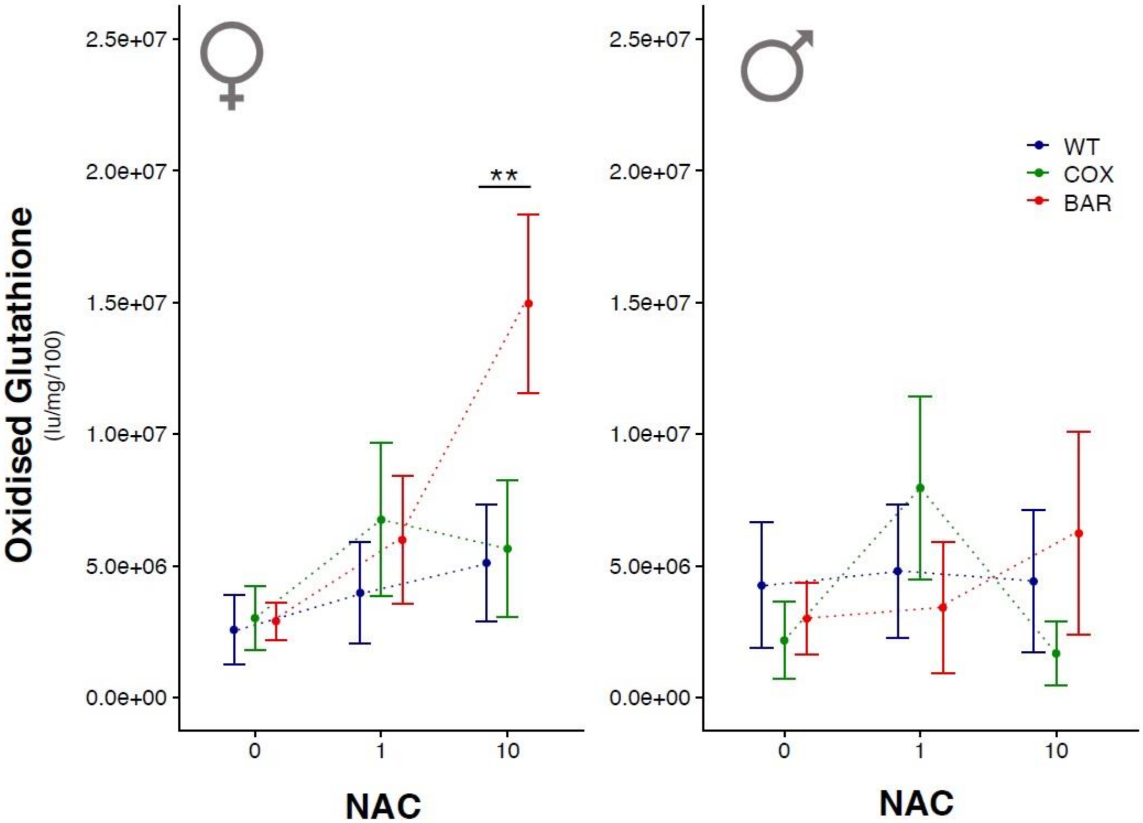


**Fig. S8 Glutathione redox state in pooled tissues for females and males**
Ratio of oxidised to reduced glutathione in pooled tissues for each mitonuclear genotype in females (left panel) and males (right panel).

#### Table S1 Contribution of each variable to a Principal component axis.

| | PC1 | PC2 |
|---|---|---|
| O2 - MPP | 0.2427903 | 0.2053067 |
| O2 - ADP | 0.2872973 | 0.2387222 |
| O2 - Succinate | 0.2902094 | 0.2412016 |
| O2 - G3P | 0.3225364 | 0.1936492 |
| O2 - FCCP | 0.32311 | 0.1945573 |
| O2 - CI | 0.3229326 | 0.1719994 |
| O2 - CII.III | 0.2867952 | 0.2319486 |
| H2O2 - MPP | 0.1597458 | -0.3691536 |
| H2O2 - ADP | 0.1625387 | -0.3729846 |
| H2O2 - Succinate | 0.1750272 | -0.3771018 |
| H2O2 - G3P | 0.2730211 | -0.2739907 |
| H2O2 - FCCP | 0.2091873 | -0.338022 |
| H2O2 - Rotenone | 0.2698082 | -0.2543651 |
| H2O2 - AntA | 0.3250485 | -0.1109828 |

#### Table S2 ANOVA results for metabolic traits across both tissues.

**Thorax**

| | O2 Flux | | Complex I Contribution | | Control Ratio | | ROS flux | |
|---|---|---|---|---|---|---|---|---|
| | F value | p-value | F value | p-value | F value | p-value | F value | p-value |
| mito | 3.2287 | **0.04423** | 3.3244 | **0.040435** | 4.3082 | **0.01637** | 0.03 | 0.97048 |
| nac | 4.7432 | **0.011** | 1.9953 | 0.141939 | 0.2871 | 7.51E-01 | 0.6483 | 5.25E-01 |
| sex | 26.6778 | **< 0.001** | 11.278 | **0.001151** | 24.2196 | **< 0.001** | 6.0794 | **0.01557** |
| mito×nac | 1.0818 | 0.37035 | 0.5254 | 0.717281 | 1.8414 | 0.12775 | 0.6998 | 0.59411 |
| mito×sex | 2.8588 | 0.06255 | 2.9546 | 0.05717 | 1.8814 | 0.1583 | 0.2804 | 0.75617 |
| nac×sex | 1.8583 | 0.16187 | 1.0958 | 0.338689 | 0.6623 | 0.51815 | 1.0375 | 0.35855 |
| mito×nac×sex | 0.2509 | 0.90842 | 0.2991 | 0.877831 | 0.1842 | 0.94609 | 0.5742 | 0.68202 |

**Reproductive Tissues**

| | O2 Flux | | Complex I Contribution | | Control Ratio | | ROS flux | |
|---|---|---|---|---|---|---|---|---|
| | F value | p-value | F value | p-value | F value | p-value | F value | p-value |
| mito | 1.9632 | 0.1457 | 8.46 | **< 0.001** | 2.0803 | 0.13018 | 0.5669 | 0.56904 |
| nac | 1.3705 | 0.2587 | 21.9751 | **< 0.001** | 0.2146 | 8.07E-01 | 0.3761 | 6.88E-01 |
| sex | 107.0622 | **< 0.001** | 37.1485 | **< 0.001** | 86.7692 | **< 0.001** | 103.6696 | **< 0.001** |
| mito×nac | 1.1844 | 0.3223 | 2.3047 | 0.0633605 | 0.451 | 0.77145 | 1.0098 | 0.40603 |
| mito×sex | 0.0398 | 0.961 | 2.9253 | 0.0581652 | 0.4511 | 0.63819 | 3.557 | **0.03212** |
| nac×sex | 12.4812 | **< 0.001** | 2.4115 | 0.0947807 | 3.2795 | **0.04169** | 6.2105 | **0.00285** |
| mito×nac×sex | 0.0484 | 0.9955 | 1.9318 | 0.1107977 | 0.5642 | 0.68917 | 1.2718 | 0.286 |
